# Blue and green food webs respond differently to elevation and land use

**DOI:** 10.1101/2021.12.22.473785

**Authors:** Hsi-Cheng Ho, Jakob Brodersen, Martin M. Gossner, Catherine H. Graham, Silvana Kaeser, Merin Reji Chacko, Ole Seehausen, Niklaus E. Zimmermann, Loïc Pellissier, Florian Altermatt

**Author notes:** **Corresponding authors:** Hsi-Cheng Ho, Florian Altermatt. **Statement of author contribution and competing interests:** FA and LP, together with JB, MG, CH, OS and NZ, developed the idea and secured the funding. HH, together with FA, SK and MRC, compiled the data. HH conducted the analyses and drafted the manuscript with substantial inputs by LP and FA. All authors contributed to subsequent revisions of the manuscript. The authors declare that they have no competing interest. **Data accessibility statement:** Taxa occurrence data and GIS environmental data can be accessed via contacting the respective authorities in charge (see *Methods*). The metaweb and local food-web data, as well as the R codes that reproduce all analyses and figures in this study, will be made available to the public upon publication in a peer-reviewed journal.

## Abstract

While aquatic (blue) and terrestrial (green) food webs are parts of the same landscape, it remains unclear whether they respond similarly to shared environmental gradients. We use empirical community data from hundreds of sites across Switzerland, and show that blue and green food webs have different structural and ecological properties along elevation as a temperature proxy, and among various land-use types. Specifically, in green food webs, their modular structure increases with elevation and the overlap of consumers’ diet niche decreases, while the opposite pattern is observed in blue food webs. Such differences between blue and green food webs are particularly pronounced in farmland-dominated habitats, indicating that anthropogenic habitat modification moderates the climatic effects on food webs but differently in blue versus green systems. These findings indicate general structural differences between blue and green food webs and suggest their potential divergent future alterations through land use or climatic changes.

## Introduction

Climate and land-use types, as well as respective anthropogenic modifications thereof, are key factors that are defining the state and change in species richness, composition, and dynamics of ecological communities [1—3]. However, communities are not just arithmetic sums of species, but groups of interacting species that form ecological networks, which underpin the structure of biodiversity and ecological functions [4, 5]. Therefore, understanding how ecological networks respond to these environmental factors is crucial for developing effective management and conservation strategies, especially in this era of global change [4, 6, 7]. Recent studies have investigated the structural changes in ecological networks along environmental gradients, as well as their effects on species persistence or ecosystem functioning [8—10]. However, investigations of network variation mostly targeted the interactions between two functionally distinctive taxonomic groups, e.g., bipartite herbivore-plant or host-parasitoid networks [11—13], or were restricted to simplified experimental systems [14]. Such partiality is likely due to methodological constraints, as measuring interactions in a standardised and comparable way across a broad range of taxonomic and functional groups is difficult. Consequently, although consumer-resource trophic interaction is arguably one of the most fundamental types of biological interaction [15, 16], our understanding of the real-world unipartite food webs formed by multiple taxonomic groups—and particularly, their responses to varying environmental conditions—remains very limited (see [17] for rare examples). Yet, such understanding is crucially needed in the contemporary biodiversity crisis affecting both aquatic and terrestrial biomes.

Aquatic (blue) and terrestrial (green) food webs have mostly been studied separately, but both are fundamental parts of the same landscape. General differences have been highlighted, where blue food webs tend to accommodate long food chains [18, 19] and have marked large-eat-small body-size relationships between consumers and resources [20, 21], thus often exhibiting a nested structure [20]. In contrast, green food webs have comparatively short food chains [18, 19] with less-prominent body-size relationships [22], and generally exhibit a modular (compartmentalised) structure [22, 23]. Despite knowing these structural differences, the pronounced separation between aquatic and terrestrial ecological research has prevented a systematic comparison between the two types of food webs along environmental gradients. Recent datasets have allowed blue-green comparisons at a global scale, e.g., among sites around the world [24], but it is still unclear whether blue and green food webs respond similarly or differently to shared environmental gradients at a landscape scale, where communities share a regional species pool. This knowledge is critically needed, as biodiversity conservation strategies for both systems need to be well-aligned to avoid lopsided management [25] given that blue and green communities can be regulated by the same environmental drivers [26].

To study spatial variation in multi-trophic food webs, metawebs that integrate the ensemble of knowledge of interactions and coupled with species co-occurrence can provide an efficient tool [27]. A metaweb is a representation of the regional food web integrating the knowledge of trophic interactions among species that are present in the target region [9, 28]. If one assumes that every consumer species has a fixed ability to feed on certain resources (i.e., a fixed potential diet set) across its geographical range, which implicitly embraces the concept that interactions are driven by matching functional characteristics and that the latter are fixed at species level [28, 29], these established trophic relationships in the metaweb allow an inference of local food webs in combination with local community data [30] (see illustration in Fig. 1). This approach can guide the identification of structural differences between food webs that result from compositional differences between local communities, and thus allows investigating how food webs may change along environmental or geographical gradients. The metaweb approach is especially adequate for comparing blue and green food webs, as the two are inherently composed of very different taxa, and thus must be standardised for an adequate inference across both systems, for example based on species co-occurrence data.

**Figure 1.**
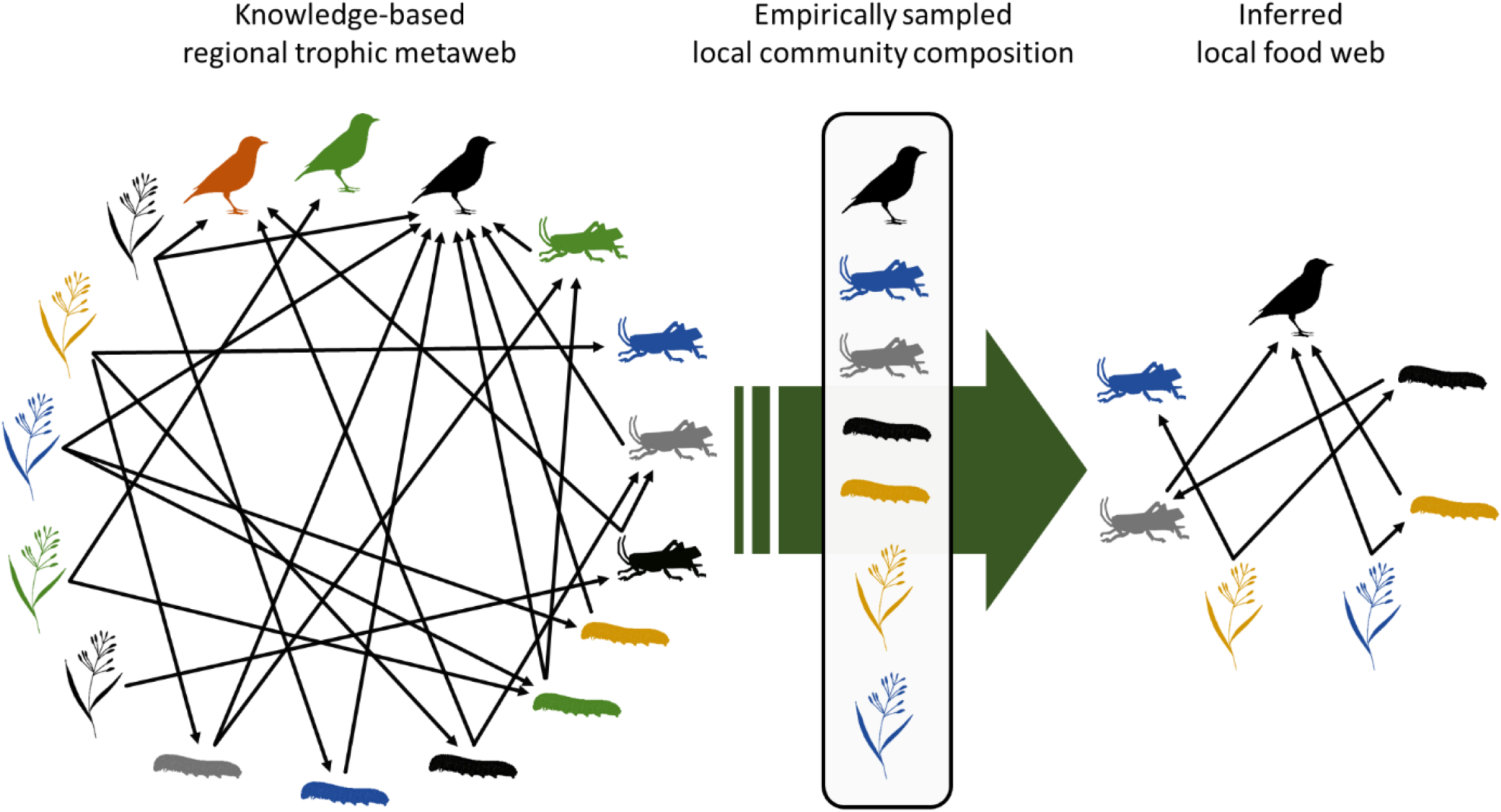
An illustration of the metaweb approach using dummy terrestrial species. Different silhouette shapes and colours represent different species, and black arrows indicate trophic relationships pointing from the resource to the consumer. Trophic relationships established in the metaweb are based on the literature and expert knowledge (left panel). Extracting trophic relationships among species that co-occur locally according to empirical survey data (middle panel) allows the inferential assembly of local food web (right panel).

Using data from highly systematic and representative species diversity monitoring schemes across freshwater and terrestrial biomes in Switzerland, which encompass the occurrence of aquatic invertebrates and fishes, as well as terrestrial plants, butterflies, grasshoppers, and birds over a 42,000 km^2^ area, we ask (i) how are food webs influenced by key environmental drivers, and (ii) do blue and green food webs respond differently to these drivers? To answer these questions, we examined the structure and expected diet niche overlap in both blue and green multi-trophic food webs inferred from a trophic metaweb. Specifically, we compared food webs between the two systems, along elevation, and across major land-use types (i.e., forests, scrubs, open spaces, farmlands and urban areas; see *Methods*). Food-web patterns discovered along elevation can be interpreted as a spatial proxy of their responses to temperature and its change, likewise patterns among land-use types as responses to habitat turnover driven by climate and/or anthropogenic activities.

## Results

We inferentially assembled local food webs for a total of 462 terrestrial (green) and 465 aquatic (blue) sites, which representatively covered the area of Switzerland and spanned an elevational range from 249 m to 2834 m a.s.l. (Fig. 2). With increasing elevation, different land-use types dominated the vicinity of local sites, and their elevational turnovers were consistently covered by the blue and green sites (Fig. 2). The inferred food webs included distinct taxa mostly at the species level of overall 2,016 plant, 191 butterfly (focused on trophic interactions of larvae), 109 grasshopper, 155 bird, 248 stream invertebrate, and 78 stream fish taxa (henceforth “focal groups”; see *Methods* for details). The blue and green food webs differed in their structural properties (i.e., number of nodes, connectance, nestedness, modularity), illustrated by the principal component analysis (along a PC1 axis explaining 70% of the variance, Fig. 3A). Across all 927 food webs, the blue food webs were in general smaller (median number of nodes in blue: 35; in green: 437), more connected (median connectance in blue: 0.25; in green: 0.06), and less modular (median modularity in blue: 0.03; in green: 0.20) than the green ones (Fig. 3A).

**Figure 2.**
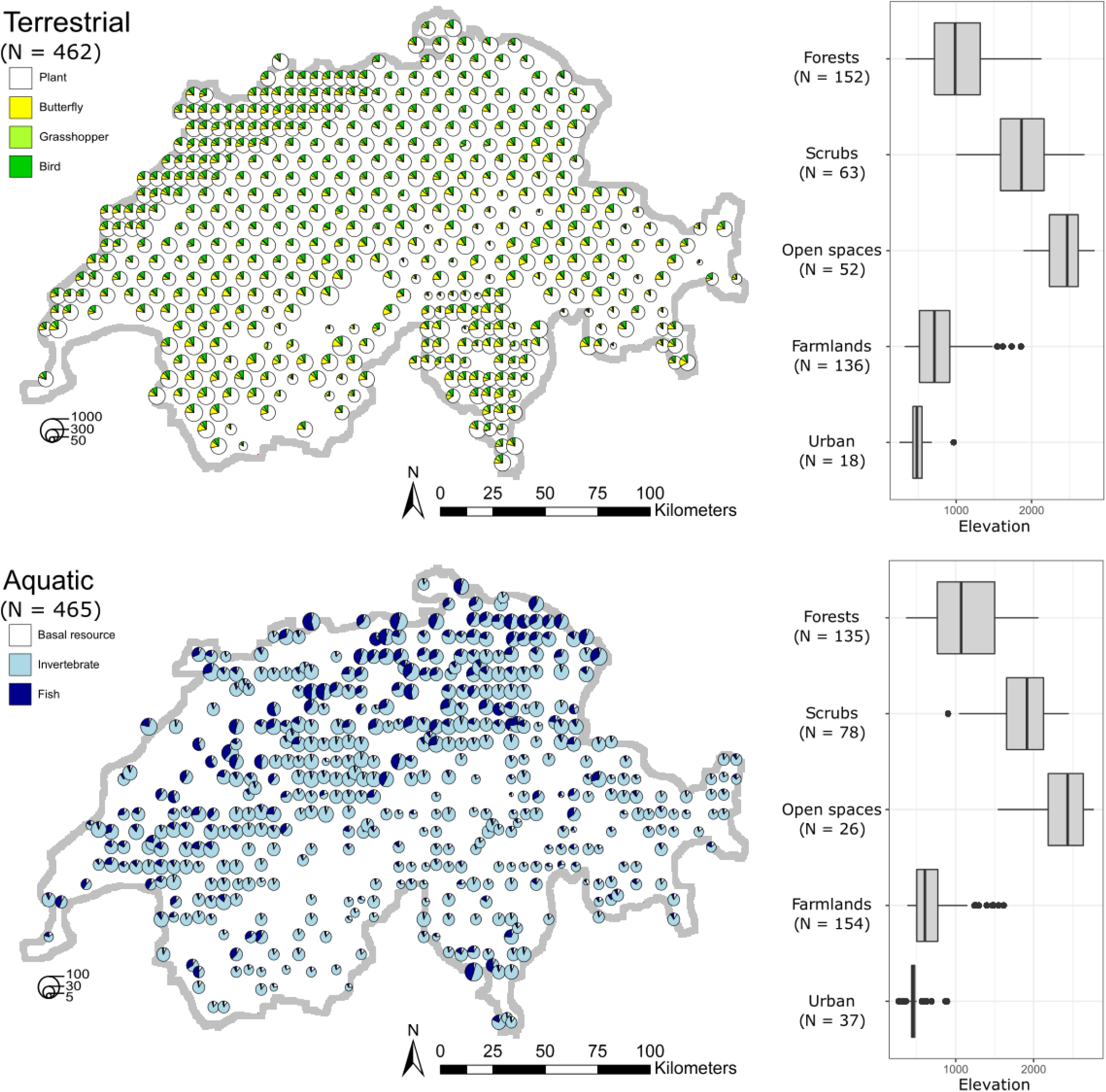
Terrestrial (top panel) and aquatic (bottom panel) food webs assembled in this study. The food webs’ locations across Switzerland are based on a stratified, randomised raster approach, and thus representative for the landscape. The pie charts give the food webs’ sizes (number of nodes, as the size of pie charts) and focal-group composition (colours). Note that the three assumptive mega nodes: plant, plankton, and detritus, served as basal resource in all aquatic food webs (*Methods*). The boxplots show, at where these food webs locate, how different dominant land-use types distribute along elevation.

**Figure 3.**
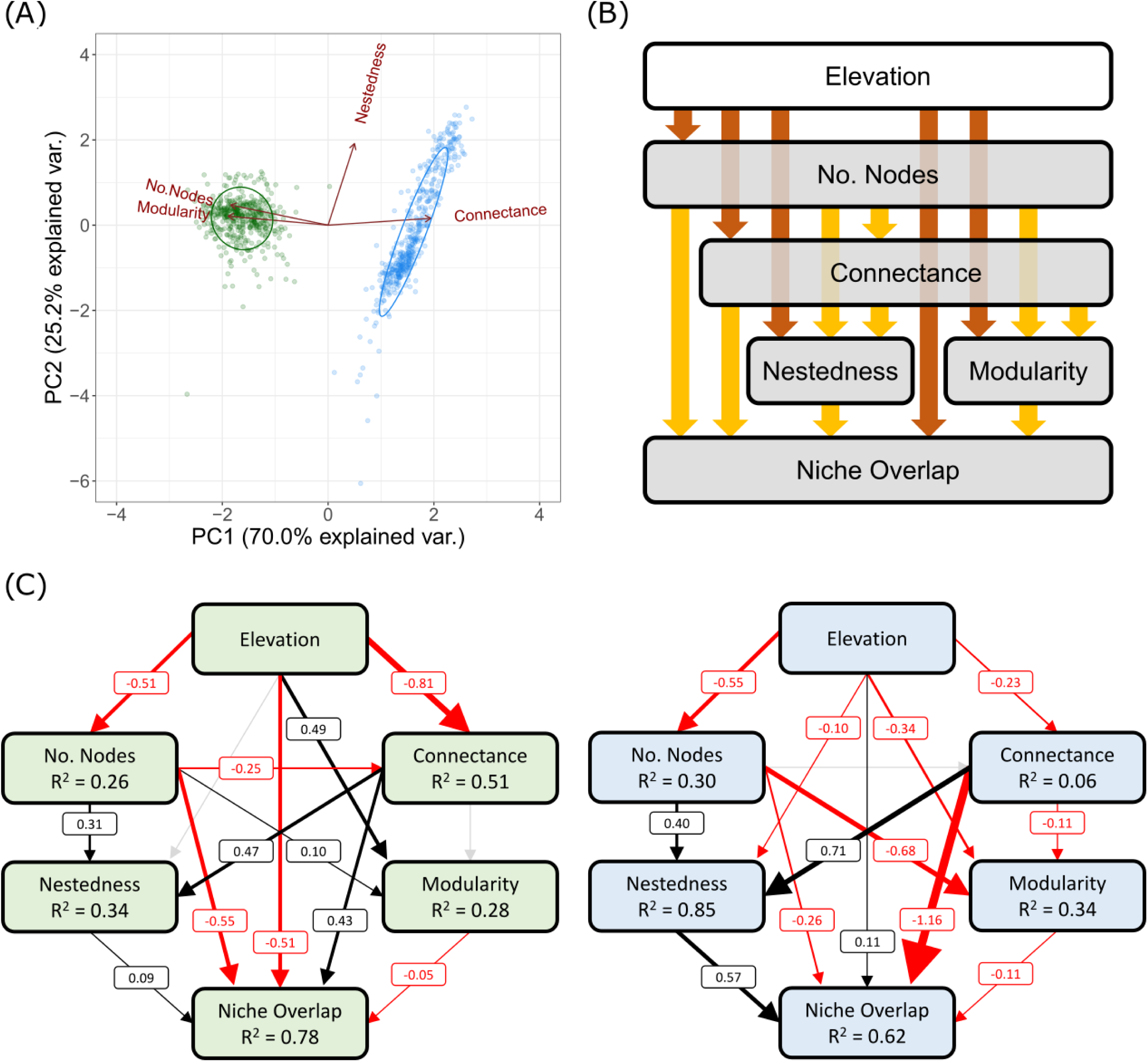
(A) The principal component analysis reveals distinct structures between the terrestrial (green) and aquatic (blue) food webs, indicated by dot colours. (B) The potential causal relationships between elevation and food-web properties (brown arrows), as well as among food-web properties themselves (yellow arrows), for the piecewise structural equation modelling (SEM) analysis. (C) The outcomes of piecewise SEM standardised coefficients and R-squared of the green and blue food webs (indicated by block colours), respectively. Positive paths in black, negative in red, and non-significant in grey. A path’s width is proportional to its size of standardised coefficient. For additional SEM results, see Figs. S1 & S2.

By considering the mutual dependence among food-web metrics using piecewise structural equation modelling (SEM) analyses, we found elevation to be significantly associated with most of the structural and ecological (i.e., diet niche overlap) properties in both blue and green food webs, but with contrasting relationships (Figs. 3B & 3C). Regarding direct effects, elevation positively influenced modularity (standardised coefficient: 0.49) and negatively influenced niche overlap (−0.51) in green food webs, while the opposite (−0.34 and 0.11, respectively) was observed in blue ones (Fig. 3C). Further SEM analyses separated by land-use types with subsetted food webs detected the same set of contrasting relationships between the two systems particularly in farmlands (Supplementary Information Fig. S1). General linear model analyses further supported that elevation and local dominant land-use type are both significant drivers of food-web properties (Table S2). Taken together, the results suggest that blue and green food webs change differently in their structural and ecological properties along elevation, resulting from each of their elevational community composition turnover, and such blue-green differences were most obvious in farmlands.

Next, to investigate potential nonlinear elevational patterns, we analysed food-web properties along elevation with generalised additive models considering all 972 food webs. In addition, to identify potential different responses among land-use types, we also conducted similar analyses with linear models comparing regression slopes of food webs subsetted by land-use types. Regarding the number of nodes (i.e., species richness), the green food webs showed an abrupt changing pattern with elevation, as the number of nodes increased with elevation until about 1500–2000 m a.s.l. but decreased thereafter (Fig. 4). This abrupt change coincides with the tree-line effect on community composition [31, 32]. The blue food webs, conversely, showed a consistent linear decrease with elevation. Further blue-green slope comparisons by land-use types revealed that the main qualitative difference between the patterns in the two systems (which occurred below 1500 m a.s.l.) was largely due to their different responses to elevation in farmlands (Fig. 4; Fig. S3): green food webs became larger because more plants and butterflies were added, while blue ones became smaller because more fishes were lost than the added invertebrates (Figs. S6–S11). The connectance decreased near linearly with increasing elevation in green food webs, whereas in blue ones connectance decreased relatively quickly with increasing elevation until 1000 m a.s.l. but mildly increased above 1000 m a.s.l. Blue food webs became less connected with increasing elevation due to the gradual loss of fishes, and thus the many trophic links toward the overall more generalist fishes’ diets, up to roughly 1000 m a.s.l. (Fig. S11). At above 1000 m a.s.l., fish became very rare and invertebrate the dominant group (Figs. S10 & S11). The increasing food-web connectance with increasing elevation thus reflected the combined effect of the shrinking food-web size (Fig. 4) and the replacement of specialist by generalist invertebrates (Fig. S10; *sensu* [33]).

**Figure 4.**
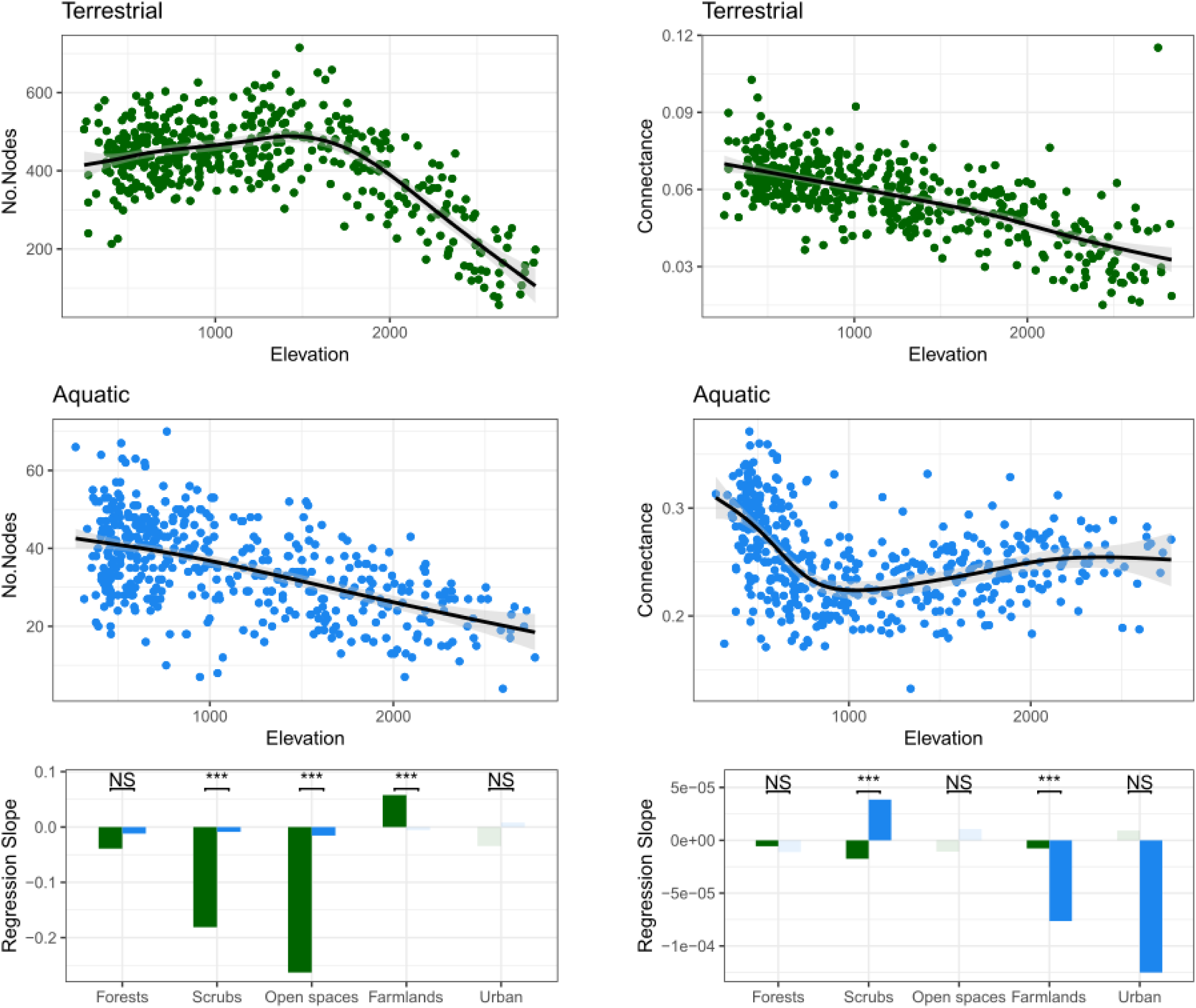
Number of nodes (left panel) and connectance (right panel) of terrestrial (green) and aquatic (blue) food webs along elevation. In the scatter plots, black lines and corresponding shades are the fitted regression and 95% CI of generalised additive models. The bottom barplots present the slope estimates from linear models testing the effects of elevation on the focal food-web property with subsetted food webs of each dominant land-use type. Solid green/blue bars indicate significant slopes, likewise faded colours non-significant ones. The significance of terrestrial versus aquatic least-squares slope comparison is indicated above the bars (*** P < 0.001, NS non-significant).

For nestedness, modularity, and diet niche overlap, we further analysed the inferred local food webs against two types of randomisation (i.e., keep-group and fully randomised webs; see *Methods*), to understand whether the observed patterns were driven by the change in food-web size and connectance, or by the change in composition of focal group or species (and thus the corresponding diet composition) along elevation. In general, green food webs were more nested and more modular (when below 2500 m a.s.l.) than their randomised counterparts (Fig. 5). Conversely, blue food webs were more nested but less modular than their randomised counterparts (Fig. 5). Both blue and green food webs showed a trend of decreasing nestedness with increasing elevation (Fig. 5). In the blue food webs, specifically, the decrease in nestedness occurred exclusively until an elevation of 1000 m a.s.l. was reached, after which nestedness plateaued (Fig. 5; Fig. S4). This rapid drop of nestedness in blue food webs again (as in connectance) matched the “fish-line” effect, that is, fish species gradually dropped out with increasing elevation, and only very few species remained above 1000 m a.s.l. (Fig. S11; see *Discussion*). Loss of fish species richness strongly influenced food-web nestedness not only because it reduced connectance (captured by fully randomised webs, Fig. 5), but also because most fish species were generalist consumers (with much broader diets than even generalist invertebrates, see Figs. S10 & S11) that tended to shape a nested structure in food webs (captured by keep-group randomised and inferred webs, Fig. 5). The green food webs, but not the blue ones, saw a mild increase of modularity with elevation (Fig. 5). At lower elevation, such an increase was due to a boost of richness with elevation in butterflies, but not grasshoppers and birds (Figs. S6–S9), leading to a higher proportion of specialist consumers within communities. However, this association between butterfly richness and elevation was inversed beyond a turning point around the tree line (Fig. S7). Thereafter, as the influences of decreasing specialist proportion (disfavouring modularity) and decreasing food-web size and connectance (favouring modularity, as shown in randomised webs in Fig. 5) could have balanced out each other, the observed modularity of green food webs became more or less unchanged (Fig. 5; Fig. S4).

**Figure 5.**
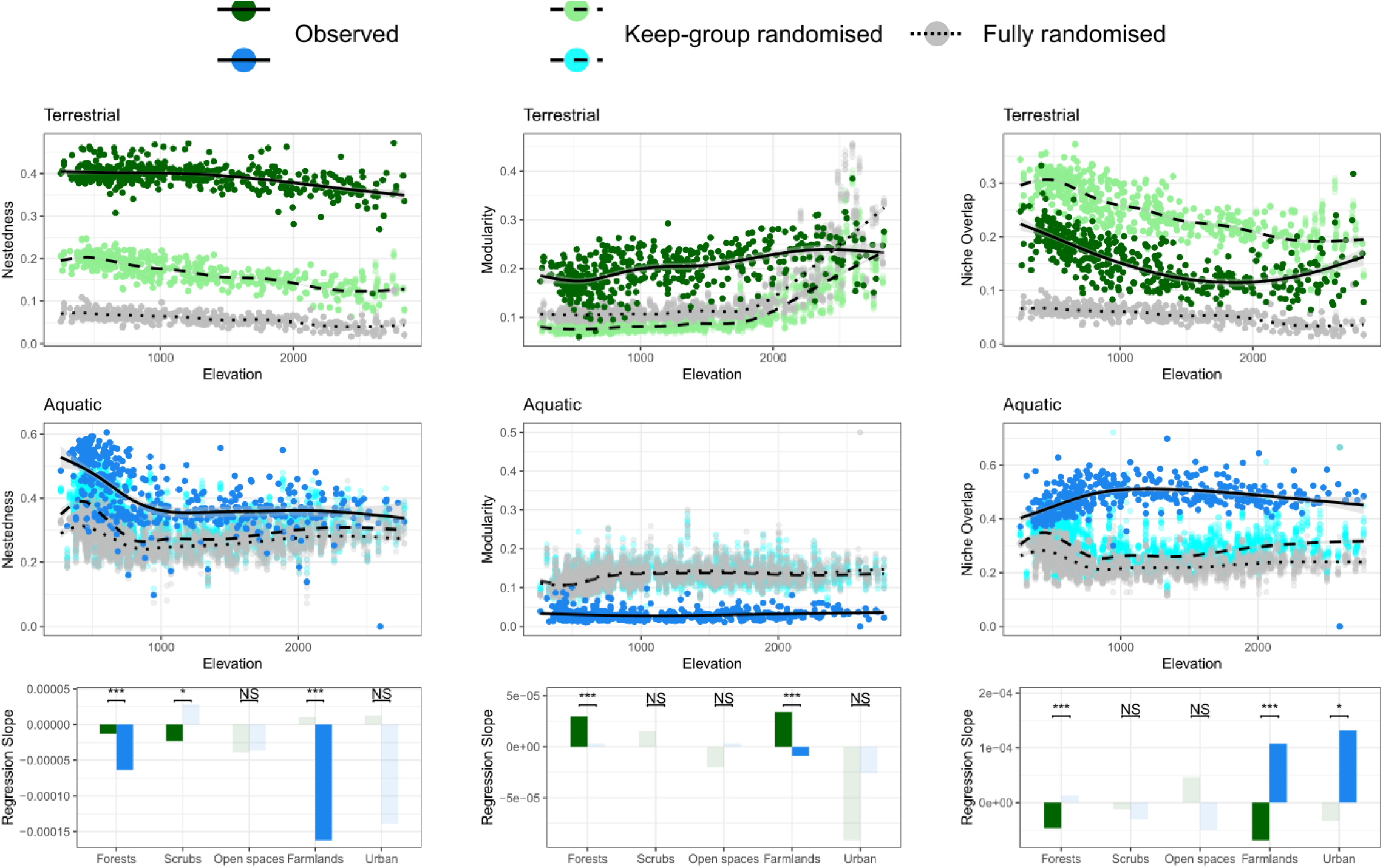
Nestedness (left panel), modularity (middle panel), and consumers’ niche overlap (right panel) of terrestrial (green) and aquatic (blue) food webs along elevation. In the scatter plots, besides the observed values from the inferred food webs (green/blue), the values from the keep-group (light green/blue) and fully (grey) randomised counterparts are presented alongside for comparison. The black lines (solid, dashed, and dotted) are the fitted regression of generalized additive models with corresponding shades the 95% CI. The bottom barplots present the slope estimates from linear models testing the effects of elevation on the focal food-web property with subsetted food webs of each dominant land-use type. Solid green/blue bars indicate significant slopes, likewise faded colours non-significant ones. The significance of terrestrial versus aquatic least-squares slope comparison is indicated above the bars (* P < 0.05, *** P < 0.001, NS non-significant).

The consumers’ diet niche overlapped more in inferred food webs than in fully randomised webs in both blue and green systems (Fig. 5), but the green food webs overlapped less while the blue ones overlapped more than their respective keep-group randomised counterparts (Fig. 5). These patterns showed that, within each of the focal consumer groups (i.e., excluding plants and aquatic basal resources), species-specific diets are more differentiated among terrestrial consumers while more overlapped among aquatic ones, compared to expectation by chance. Both types of food webs had nonlinear patterns of niche overlap along elevation, but in different ways: the green food webs saw a decrease until about 1500–2000 m a.s.l., and switched to an increase thereafter (Fig. 5), again reflecting the effects of the tree line and the change in the richness of specialist butterflies (Fig. S7). Contrastingly, the blue food webs saw an increase in niche overlap until about 1000 m a.s.l. then remained roughly constant thereafter (Fig. 5). This increasing niche overlap together with the decreasing nestedness along elevation up to 1000 m a.s.l. reflected that, as fish species richness decreased along elevation, it was those relatively-generalist invertebrates sharing alike diets that became the majority in stream communities (Fig. S10) and determined food-web structure.

For all five food-web metrics analysed against elevation, we found significant differences in blue versus green regression slopes in farmlands (Figs. 4 & 5; Figs. S3 & S4). There specifically, the blue and green slopes were both statistically significant and having opposite signs in two metrics, such that modularity was positively and diet niche overlap negatively associated with elevation in green food webs, and the inverse in blue ones (Figs. 4 & 5; Figs. S3 & S4). These were the most pronounced blue-green differences we detected among all land-use types. In accordance with the findings from our SEM analyses by land-use type, in general, it was especially in farmlands where the blue and green food webs exhibited qualitatively different responses to elevation.

## Discussion

Examining ecological communities from the perspective of not only the richness of species, but also the interaction network they form, can inform management of biodiversity and ecosystem functioning [4, 6]. We showed that local food webs exhibit variation in their structural and ecological properties corresponding to a change in elevation and variation in surrounding land-use types. Specifically, terrestrial (green) and aquatic freshwater (blue) food webs responded differently to these environmental factors. This suggests that the same environmental change can influence terrestrial and aquatic taxa differently, thereby imposing complicated overall impacts on the biodiversity and functions of local ecosystems (*sensu* [34]). As environmental gradients are reshaped by global changes, the complex responses of food webs challenge the development of effective management and conservation strategies.

Using nearly a thousand inferred local food webs representatively covering Switzerland, we saw clear and gradual changes in almost all key food-web metrics with elevation in both blue and green systems. In many cases, the overall pattern was nonlinear. Some of such nonlinear food-web responses along elevation were driven by bio-geographical boundaries, e.g., the tree line (around 1500–2000 m a.s.l. across our local sites, see Fig. 2) in green food webs, while some were associated with food-web responses specific to land-use types in combination with a land cover turnover by elevation (Fig 2; Figs. S3–S5). The elevation gradient is often taken as a proxy of climatic change in ecological research (e.g., [35]), and important climatic variables, such as temperature and precipitation, change consistently along elevation. Thereby, the herein considered elevation gradient captures among others effects of temperature on food webs (*Methods*; Table S3). In terrestrial systems, our focal consumers, i.e., butterflies, grasshoppers, and birds, are mostly highly mobile. Their elevational distributions are thus expected to be influenced not so much by topographical constraints as by their physiological constraints in relation to temperature and bottom-up reliance on temperature-shaped vegetation [36—38]. Therefore, the green food-web patterns we observed along elevation likely reflected the effects of elevational temperature variation on species distribution and thus the composition of communities. In contrast, in stream systems, the existence of many taxa is also constrained by the steepness (thus flow speed) or width of the stream, which both covariate with elevation [39, 40]. For instance, only very few fish species, mostly brown trout, are present above 1000 m a.s.l. (i.e., the “fish line”, Fig. S11) because the structure and hydrology of streams does not favour many fishes at these elevations. Indeed, we saw a fish-line effect in several detected elevational patterns of our blue food webs (Figs. 4 & 5). The temperature effects are nonetheless still identifiable, as the observed patterns below the fish line were broadly in line with temperature-driven ones reported in the literature, such that higher temperature favours the existence of fishes in streams [41] (Fig. S11) and higher food-web connectance [42] (Fig. 4). Overall, the blue food-web patterns along elevation are likely shaped simultaneously by both temperature and topographical effects.

The nested and non-modular structure detected in the blue food webs (Fig. 5) well-echoes the reports in the literature [43, 44], reflecting that aquatic consumers are usually broad feeders with diets mostly constrained by body size (or say gape size) instead of other feeding traits [45, 46]. In contrast, the relatively high modularity observed in the green food webs (Fig. 5) suggests a relatively high proportion of specialised consumer-resource pairs within communities [47] (Fig. S7). This is further supported by the fact that green food webs had lower while the blue ones had higher diet niche overlap than their respective keep-group randomised counterparts (Fig. 5). It is an inherent property of aquatic and terrestrial food webs, respectively, that the latter contain a very large part of highly specialised co-evolved interaction partners (i.e., plant-insect co-evolution), while the former are more constrained by resource availability [46]. Therefore, diet niche differentiation versus overlap may not be as important a constraining factor for species coexistence in stream communities as in terrestrial ones. Interestingly, in terms of diet specialisation and differentiation, elevation had significant yet opposite influences on blue and green food webs, especially in farmlands (Figs. 3C & 5; Figs. S1 & S4). With increasing elevation and thus decreasing temperature, green food webs were increasingly composed by more-specialist butterflies and birds whose diets were more differentiated, whereas blue food webs by more-generalist fishes and invertebrates whose diets were more overlapped (Figs. S6–S11). These findings thus suggest different elevation-dependent strategies for managing local biodiversity between the two systems. Toward higher elevation, we should emphasise more on keeping diverse resources in green communities to support the living of specialists, meanwhile stabilising resource quantities in blue communities, given the high diet niche overlap can impose excessive competition among consumers when the shared resources become rare [48, 49]. In addition, with a global-change prospect, climatic warming can threaten biodiversity via imposing physiochemical conditions that no longer suits local populations [50], or triggering biological range shift where new biological interactions emerge [51]. According to our findings, terrestrial communities may be especially vulnerable to the latter mechanism, such that warming can spark an elevational upshift of generalist species into regions originally dominated by specialists, provoking predation or resource competition pressures and thus increasing threat to the latter.

Anthropogenic land use has been shown to be a major driver of the richness and community composition of different taxonomic and trophic groups (e.g., [34, 52]), but rarely associated with food-web properties (e.g., [12]), and here for the first time is compared between blue and green systems. Our focal land-use types are defined based on the vegetation or human modifications on land, which often also cascade to aquatic systems, but are not describing the streams themselves that are embedded into the terrestrial matrix. Nonetheless, we detect significant land-type effects not only in green but also in blue food webs (Figs. 4 & 5; Figs. S3 & S4). There are various interchanges between terrestrial and aquatic systems that may provide channels for potential cross-ecosystem spill-over effects on community composition [53]. For instance, leaves and terrestrial organisms become the inputs of organic matter to water bodies, and many aquatic insects at their late life stages emerge into terrestrial communities [54, 55]. However, in our case, the comparison among blue food webs with controlled elevation revealed no obvious difference between forests and farmlands (Fig. S5), indicating that the detected land-use effects in blue food webs reflect mostly their elevational responses in combination with changes in land-use type along elevation, rather than spill-over effects from land. Regarding the green food webs, the land-use type *per se* showed strong effects. Specifically, green food webs in farmlands tended to be smaller and have higher overlap in diet niche than those in forests at low elevation, but not at higher elevation (Fig. S5). Compared to natural conditions, agriculture generally simplifies local vegetation into a few selected crops and exerts frequent perturbations (e.g., harvesting) on the habitat, leading to conditions that favour generalists over specialists [56]. Indeed, at low-elevation farmlands where agricultural intensity is relatively high, the terrestrial communities fostered fewer plants, and fewer but more-generalist butterflies, than at either high-elevation farmlands or low-elevation forests (Figs. S6 & S7). The different elevational food-web patterns we observed in forest versus farmland green food webs suggest that anthropogenic land use and climate change can have interactive ecological impacts, such that the warming effect on food webs can be moderated by human-caused habitat modifications and its intensity.

In conclusion, our large-scale analysis of food-web structure and change in blue and green systems provides the first evidence that blue and green food webs within the same landscape respond differently (in terms of major network metrics) to elevation and land use. The food-web patterns featured in this study emerge spatially from community compositional differences, which supposedly reflect the outcome of both evolutionary and ecological processes. By making analogies between elevation and climate change, our findings provide not only a broad depiction of both blue and green food webs with their current status across the landscape, but also visionary implications with their potential future change. Such understandings could become fundamental knowledge when managing local biodiversity and ecosystem functioning, especially in places where blue and green communities coexist and is vulnerable to anthropogenic modifications.

## Methods

### Overview

We compiled systematically sampled empirical taxa occurrence across the landscape, and inferentially assembled respective blue and green local food webs combining these data with a metaweb approach. We quantified key properties of the inferred food webs, then analysed with GIS-derived environmental information how focal food-web metrics change along elevation and among different land-use types in blue versus green systems. Details are given below.

### Assemble food webs using a metaweb approach

We applied a metaweb method to obtain the composition and structure of multiple local food webs across a landscape spatial scale [9]. A metaweb is an accumulation of all known interactions (here, trophic relationships) among focal taxa. By assuming that any interaction in the metaweb will realise if the interacting taxa co-occur, the metaweb approach allows an inference of local food webs if taxa occurrence is known. The assumption of fixed diets may lead to an over-estimation of the locally realised trophic links [29], as it essentially ignores the possible intraspecific diet variation caused by resource availability [57, 58], predation risk [59], temperature [60], ontogenetic shift [61], or other genetic and environmental sources [62]. Therefore, the food webs we inferred systematically using this method capture trophic relationships driven by community composition (species presence versus absence) but not the above-mentioned processes.

We compiled taxa occurrence of four terrestrial and two aquatic broad taxonomic groups (“focal groups”) to assemble local green and blue communities, respectively and independently, based on the well-resolved data available. Each focal group referred to a distinct taxonomic group, and the within-and among-group trophic relationships captured most of the realised interactions. These focal groups were plants, butterflies, grasshoppers, and birds in the green biome, and stream invertebrates and fishes in the blue biome. Notably, with “butterflies” we refer to their larval stage and accordingly their mostly-herbivorous trophic interactions throughout this study. The occurrence data of these focal groups were compiled from highly standardised multiple-year empirical surveys of various authorities, all conducted by trained biologists with fixed protocols. The information across sites should thus be representative and can be up-scaled to the landscape. The occurrences of plants, butterflies, birds, and stream invertebrates were from the *Biodiversity Monitoring Switzerland* programme (BDM Coordination Office [63]) managed by the Swiss Federal Office for the Environment (BAFU/FOEN). The occurrences of grasshoppers and fishes were from the Swiss database on faunistic records, *info fauna* (CSCF), where we further complemented fish occurrence from the data of *Progetto Fiumi Project* (Eawag). In terms of biological resolution, taxa were resolved to species level in most cases, while the plant and butterfly groups included some multi-species complexes. Insects of the order Ephemeroptera, Plecoptera and Trichoptera were resolved to species, while all other stream invertebrates were resolved to family level. These were each treated as a node later in our food-web assembly, and referred to as “species”, as the species within such complexes and families mostly share the same trophic role. Spatially, the occurrence datasets adopted coordinates resolved to 1×1 km^2^. The species that were recorded in the same 1×1 km^2^ grid were considered to co-occurred. We took the co-occurring four/two focal groups to form local green/blue local communities, respectively. To obtain better co-occurrence across group-specific data from different sources (e.g., *BDM* and *info fauna*), we intentionally coarsened the grasshopper and fish occurrence to 5×5 km^2^ coordinates. This is arguably a biologically acceptable approximation considering the high mobility of these two groups. Also, we only included known stream-borne fishes and dropped pure lake-borne ones to match our stream-only invertebrate occurrence data. Across all 462 green and 465 blue communities we assembled, we covered 2016 plant, 191 butterfly, 109 grasshopper, 155 bird, 248 stream invertebrate, and 78 stream fish species. Unlike the knowledge of plant occurrence in green communities, we did not have detailed occurrence information of the basal components (e.g., primary producers) in blue ones. Therefore, we assumed three mega nodes—namely plant (including all alive or dead plant materials), plankton (including zooplankton, phytoplankton, and other algae), and detritus—as the basal nodes occurring in every blue communities, without further discrimination of identities or biology within. These adding to our focal groups thus cover major taxonomic groups as well as trophic roles from producers to top consumers in both blue and green systems.

We built our metaweb based on reported trophic interactions among the focal taxa in literature, then complemented it with expert knowledge. The respective literature we referred, as well as a link to the metaweb itself, are provided in Supplementary Information Section S1. Taking the above-assembled local communities then drawing trophic links among species (nodes) according to the metaweb yielded the local food webs (illustrated in Fig. 1), representatively covering the whole Swiss area. Notably, although our understanding of trophic interactions indeed encompassed some links across the blue and green taxa (e.g., between piscivorous birds and fishes), our occurrence datasets did not present sufficient spatial grids where these taxa co-occur. We therefore did not include such links, nor assembled blue-green interconnected food webs, but the blue and green food webs separately instead. Also, we dropped isolated nodes, i.e., basal nodes without any co-occurring consumer and consumer nodes without any co-occurring resource, from the inferred food webs. These could possibly be passing-by species that were recorded but had no trophic interaction locally, or those that interact with non-focal taxa whose occurrence information was unknown to us. We thus had to exclude them to focus on evidence supported occurrences and trophic interactions. Nonetheless, across all cases, isolated nodes were rather rare (averaged less than 3% of species occurred in either blue or green communities).

### Environmental data

We acquired environmental data across Switzerland on a 1×1 km^2^ grid basis (i.e., values are averaged over the grid) from GIS databases, with which we mapped environmental conditions to the grids where we assembled food webs. These included: topographical information from *DHM25* (Swisstopo, FOT), land-cover information from *CLC* (EEA), and climate information (averaged over the decade of 2005–2015) from *CHELSA*. Among environmental variables, elevation and temperature are essentially highly correlated. In this study, we took elevation as the focal environmental gradient throughout, as after accounting for the main effects of elevation on temperature, the residual temperature was not a good predictor of the food-web metrics we looked at (see next section, and Table S3). In other words, by analysing along the elevation gradient, we already captured most of the temperature influences on food webs. Based on the labels provided by the GIS databases, we categorised the originally detailed land cover into the five major land-use types that we used in this study, namely forest, scrubland, open space, farmland, and urban area. Forest includes broad-leaved, coniferous, and mixed forests. Scrub includes bushy and herbaceous vegetation, heathlands, and natural grasslands. Open space encompasses sparsely vegetated areas, such as dunes, bare rocks, glaciers and perpetual snow. Farmland include any form of arable, pastures, and agro-forestry areas. Finally, urban area is where artificial constructions and infrastructure prevail. As each grid could contain multiple land-use types, we then defined the dominant land-use type of the grid as any of the five above that occupied more than 50% of the grid’s area. Analyses separated by land-use types with subsetted food webs (land-use-specific analyses) were based on the grids’ dominant land-use type. There were a few grids where the dominant land-use type did not belong to the focal major five, e.g., wetlands or water bodies, and a few where no single land-use type covered more than 50% of the area. Food webs of these grids were still included in the overall analyses but excluded from any land-use-specific analyses (as revealed in the difference in sample sizes between all versus land-use type subsetted food webs in Fig. 2; analyses details below).

### Food-web metrics and analyses

We quantified five metrics as the measures of the food webs’ structural and ecological properties. For the fundamental structure of the food webs, the number of nodes (“No. Nodes”) reflects the size of the web, meanwhile represents local species richness (though the few isolated nodes were excluded as above-mentioned). Connectance is the proportion of realised links among all potential ones (thus bounded 0–1), reflecting how connected the web is. We also derived holistic topological measures, namely nestedness and modularity. Nestedness of a food web, on the one hand, describes the tendency that some nodes’ narrower diets being subsets of other’s broader diets. We adopted a recently developed UNODF index [64] (bounded 0– 1) that is especially suitable for quantifying such a feature in our unipartite food webs. On the other hand, modularity (bounded 0–1 with our index) reflects the tendency of a food web to form modules, where nodes are highly connected within but only loosely connected between. Nestedness and modularity are two commonly investigated structures in ecological networks and have been considered relevant to species feeding ecology [22] and the stability of the system [65]. Finally, we measured the level of consumers’ diet niche overlap of the food webs (Horn’s index [66], bounded 0–1), which essentially depends on the arrangement of trophic relationships (thus the structure of the webs), and could have strong ecological implications as niche partitioning has been recognised to be a key mechanism that drives species coexistence [67, 68].

To first gain a glimpse of the structure of the blue and green food webs, we performed a principal component analysis (PCA; Fig. 3A) on the inferred food webs taking the four structural metrics (number of nodes, connectance, nestedness, and modularity) as the explaining variables of blue versus green system types. We then confirmed that system type, elevation, and land-use type were all important predictors of food-web metrics (whereas the residual temperature after accounting elevation effects was not) by conducting general linear model analyses, taking the former as interactive predictors while the latter response variables (Tables S2 & S3). To check how elevation influences food-web properties in blue and green systems separately, and how food-web properties depend on each other, we ran a series of piecewise structural equation modelling (SEM) [69] analyses on all inferred food webs (Figs. 3B & 3C) and on subsetted webs of each of the five major land-use types (Figs. S1 & S2). The structure of direct effects was set according to the literature [12, 65] and is illustrated in Fig. 3B. Finally, to check and visualise the exact changing patterns of food webs, we applied generalised additive models (GAMs) to reveal the relationships between food-web metrics and the whole-ranged elevation (Figs. 4 & 5), as well as a particular comparison between food webs in forests and farmlands below 1500 m a.s.l. (Fig. S5), as this elevation segment covered most of the sites belonged to these two land-use types. We also performed a series of linear models (LMs) and least-squared slope comparisons based on land-use-specific subsets of food webs (Figs. 4 & 5; Figs. S3 & S4), to investigate whether food-web elevational patterns are different among land-use types. In the GAMs analyses, specifically, we simulated two sets of randomised webs, i.e., “keep-group” and “fully”, as the null models to compare with the inferred ones [70]. Both randomisations generated ten independently simulated webs from each input inferred local food web, keeping the same number of nodes and connectance as of the latter. On the one hand, the keep-group randomisation shuffled trophic links from an input local web but only allowed them to realised fulfilling some pre-set within- and among-group relationships. That is, in green communities, birds can feed on all groups, grasshoppers on any groups but birds, while butterflies only on plants; in blue communities, fishes can feed on all groups, while invertebrates on themselves and the basal resources. These pre-set group-wide relationships captured the majority of realistic trophic interactions compiled in our metaweb. On the other hand, the fully randomised webs shuffled trophic links disregarding the biological identity of nodes. The GAMs of nestedness, modularity, and niche overlap illustrated the patterns of these randomised webs (Fig. 5). Comparing among the three types of webs, the patterns exhibited already by fully randomised webs should be those contributed by variations in web size and connectance, while the difference between keep-group and fully randomised webs by the focal-group composition of local communities, and the difference between inferred and keep-group randomised webs further by the realistic species-specific diets. In addition, we also applied the same GAMs and LMs approach to analyse node richness, as well as both realised and potential diet generality (vulnerability for plants) of each focal group (Figs. S6–S11). These analyses provided hints about the changes in community composition and species diet breadths along elevation and among land-use types, which helped explain the detected food-web responses in mechanistic ways.

All metric quantification and analyses were performed under R version 4.0.3 (R Core Team [71]). All applied packages and functions were described in Section S2, while the R scripts performing these tasks can be accessed at the online repository provided.

## Acknowledgement

We thank the ETH Board for funding through the Blue-Green Biodiversity (BGB) Initiative (BGB2020). We thank the Swiss Federal Office of the Environment (FOEN) and the Coordination Office of the Biodiversity Monitoring Switzerland (*BDM*, especially Tobias Roth) for providing plant, butterfly, and bird occurrence data from the *Schweizerische Vogelwarte* and the *BDM*, as well as the Centre Suisse de Cartographie de la Faune (CSCF) and Yves Gonseth for providing grasshopper and fish occurrence data from the *info fauna*. We thank Rosi Siber for extracting needed GIS data. We thank Raffael Ayé, Thomas Sattler, Felix Neff, Luiz Jardim De Queiroz, and Carmela Dönz for their respective assistance in compiling trophic relationships among the focal groups in this study.

## Notes

### Competing Interest Statement

The authors have declared no competing interest.

