## Supplementary information for "Blue and green food webs respond differently to elevation and land use"

*** Corresponding authors:**

**S1. The metaweb**

We here provide a meta-data table of our metaweb. The trophic interactions among focal groups were first assigned based on references, then further complemented from experts' knowledge by the authors and collaborators. The metaweb *per se* as tables of consumer-resource links can be accessed at the provided online repository. Note that cannibalistic links were not included in this study.

**Table S1**: Meta-data of the metaweb.

| **Trophic information** | **Taxonomic scale** | **References** (ordered by year) |
|---|---|---|
| Butterfly-plant (larvae-host) interactions | Species/Sp.-complex | Landolt et al. (2010)<br>Ebert (1991–2005) |
| Grasshoppers' diets | Species | Detzel (1998)<br>Ingrisch & Köhler (1998)<br>Maas et al. (2002)<br>Schlumprecht & Waeber (2003)<br>Baur et al. (2006)<br>Landolt et al. (2010)<br>Pitteloud et al. (2020) |
| Birds' diets | Species | Storchová & Hořák (2018) |
| Aquatic invertebrates feeding group assignment and trophic relationships among groups | Species/Family | Moog (1995)<br>Schmedtje & Colling (1996)<br>*AQEM expert consortium* (2002)<br>Graf et al. (2008; 2009)<br>Tachet et al. (2010)<br>*freshwaterecology.info*: Schmidt-Kloiber & Hering (2015) |
| Fishes' diets | Species/Sp.-complex | Kottelat & Freyhof (2007)<br>*FishBase*: Froese & Pauly (2010) |

**S2. Food-web metrics quantifying and analysing tools**

All metric quantification and analyses were done in R language. For quantifying food-web nestedness, we used the *unodf* function from *UNODF* package (Cantor et al. 2017). For quantifying food-web modularity, we used in combination the *graph.adjacency*, *multilevel.community*, and *modularity* functions from *igraph* package (Csardi & Nepusz 2006). For quantifying diet niche overlap of the consumers, we adopted the inbuilt Horn's index calculation of the *networklevel* function from *bipartite* package (Dormann et al. 2009).

We carried out the principal component analysis using R-base *prcomp* function, then visualised the result using the *ggbiplot* function from *ggbiplot* package (Vu 2011). The general linear model analyses were conducted using R-base *lm* and *anova* functions, while the structural equation modelling analyses using the *psem* function from *piecewiseSEM* package (Lefcheck 2016). We plotted food-web metrics against elevation, and performed the relevant generalised additive models and linear models analyses, using the ggplot2 package (Wickham 2016) and its inbuilt analysing functions. The land-use-specific linear model slope comparisons were carried out using the *lm* function and *lstrends* function of *lsmeans* package (Lenth 2016) in combination. We illustrated the location and composition of our food webs on a Swiss map (area frame) using the *geom_scatterpie* function from *scatterpie* package (Yu 2021) and function *png* from *png* package (Urbanek 2013). The *ggpubr* (Kassabara 2020), *ggpmisc* (Aphalo 2020), and *gridExtra* (Auguie 2017) packages were also applied alongside *ggplot2* for generating needed components of the plots.

### S3. Additional tables and figures

**Table S2.** General linear model analyses with observed properties of the inferred food webs as response variable, while elevation (continuous), system (2 levels: terrestrial versus aquatic), and dominant land-use type (5 levels: forests, scrubs, open spaces, farmlands, urban) as interacting predictors. Significance code: * P < 0.05, ** P < 0.01, *** P < 0.001.

Model: Response variable ~ Elevation (Ele) * System (Sys) * Dominant Land-use Type (DLT)

| **No.Node** | Df | F value | P value |
|---|---|---|---|
| Ele | 1 | 142.987 | < 2.2e-16 *** |
| Sys | 1 | 13687.882 | < 2.2e-16 *** |
| DLT | 4 | 91.803 | < 2.2e-16 *** |
| Ele * Sys | 1 | 151.173 | < 2.2e-16 *** |
| Ele * DLT | 4 | 29.586 | < 2.2e-16 *** |
| Sys * DLT | 4 | 49.350 | < 2.2e-16 *** |
| Ele * Sys * DLT | 4 | 21.416 | < 2.2e-16 *** |

| **Connectance** | Df | F value | P value |
|---|---|---|---|
| Ele | 1 | 332.531 | < 2.2e-16 *** |
| Sys | 1 | 10798.702 | < 2.2e-16 *** |
| DLT | 4 | 11.302 | 6.2e-9 *** |
| Ele * Sys | 1 | 0.787 | 0.375 |
| Ele * DLT | 4 | 9.253 | 2.6e-7 *** |
| Sys * DLT | 4 | 10.468 | 2.8e-8 *** |
| Ele * Sys * DLT | 4 | 15.259 | 4.8e-12 *** |

| **Nestedness** | Df | F value | P value |
|---|---|---|---|
| Ele | 1 | 232.542 | < 2.2e-16 *** |
| Sys | 1 | 3.965 | 0.047 * |
| DLT | 4 | 11.558 | 3.9e-9 *** |
| Ele * Sys | 1 | 59.518 | 3.5e-14 *** |
| Ele * DLT | 4 | 4.097 | 0.003 ** |
| Sys * DLT | 4 | 10.191 | 4.7e-8 *** |
| Ele * Sys * DLT | 4 | 11.517 | 4.2e-9 *** |

| **Modularity** | Df | F value | P value |
|---|---|---|---|
| Ele | 1 | 417.484 | < 2.2e-16 *** |
| Sys | 1 | 8849.818 | < 2.2e-16 *** |
| DLT | 4 | 7.814 | 3.5e-6 *** |
| Ele * Sys | 1 | 127.811 | < 2.2e-16 *** |
| Ele * DLT | 4 | 2.432 | 0.046 * |
| Sys * DLT | 4 | 10.401 | 3.2e-8 *** |
| Ele * Sys * DLT | 4 | 2.791 | 0.025 * |

| **Niche Overlap** | Df | F value | P value |
|---|---|---|---|
| Ele | 1 | 159.672 | < 2.2e-16 *** |
| Sys | 1 | 12488.197 | < 2.2e-16 *** |
| DLT | 4 | 3.470 | 0.008 ** |
| Ele * Sys | 1 | 163.613 | < 2.2e-16 *** |
| Ele * DLT | 4 | 3.586 | 0.007 ** |
| Sys * DLT | 4 | 27.501 | < 2.2e-16 *** |
| Ele * Sys * DLT | 4 | 20.263 | 6.1e-16 *** |

**Table S3.** General linear model analyses with observed properties of the inferred food webs as response variable, while elevation (continuous) and the residual temperature (continuous) as independent predictors. The residual temperature are derived from removing the linear regression main effects of elevation on temperature. We note that elevation is the better predictor over residual temperature throughout. Significance code: * P < 0.05, ** P < 0.01, *** P < 0.001.

| Model: Response variable ~ Elevation (Ele) + Residual Temperature (ResTem) | | | | | | | |
|---|---|---|---|---|---|---|---|
| **No.Node** | Df | F value | P value | **Connectance** | Df | F value | P value |
| Ele | 1 | 7.870 | 0.005 ** | Ele | 1 | 23.923 | 1.2e-6 *** |
| ResTem | 1 | 1.758 | 0.185 | ResTem | 1 | 2.023 | 0.155 |
| **Nestedness** | Df | F value | P value | **Modularity** | Df | F value | P value |
| Ele | 1 | 189.292 | < 2.2e-16 *** | Ele | 1 | 36.007 | 2.9e-9 *** |
| ResTem | 1 | 1.776 | 0.183 | ResTem | 1 | 6.064 | 0.014 * |
| **Niche Overlap** | Df | F value | P value | | | | |
| Ele | 1 | 9.920 | 0.002 ** | | | | |
| ResTem | 1 | 3.307 | 0.069 | | | | |

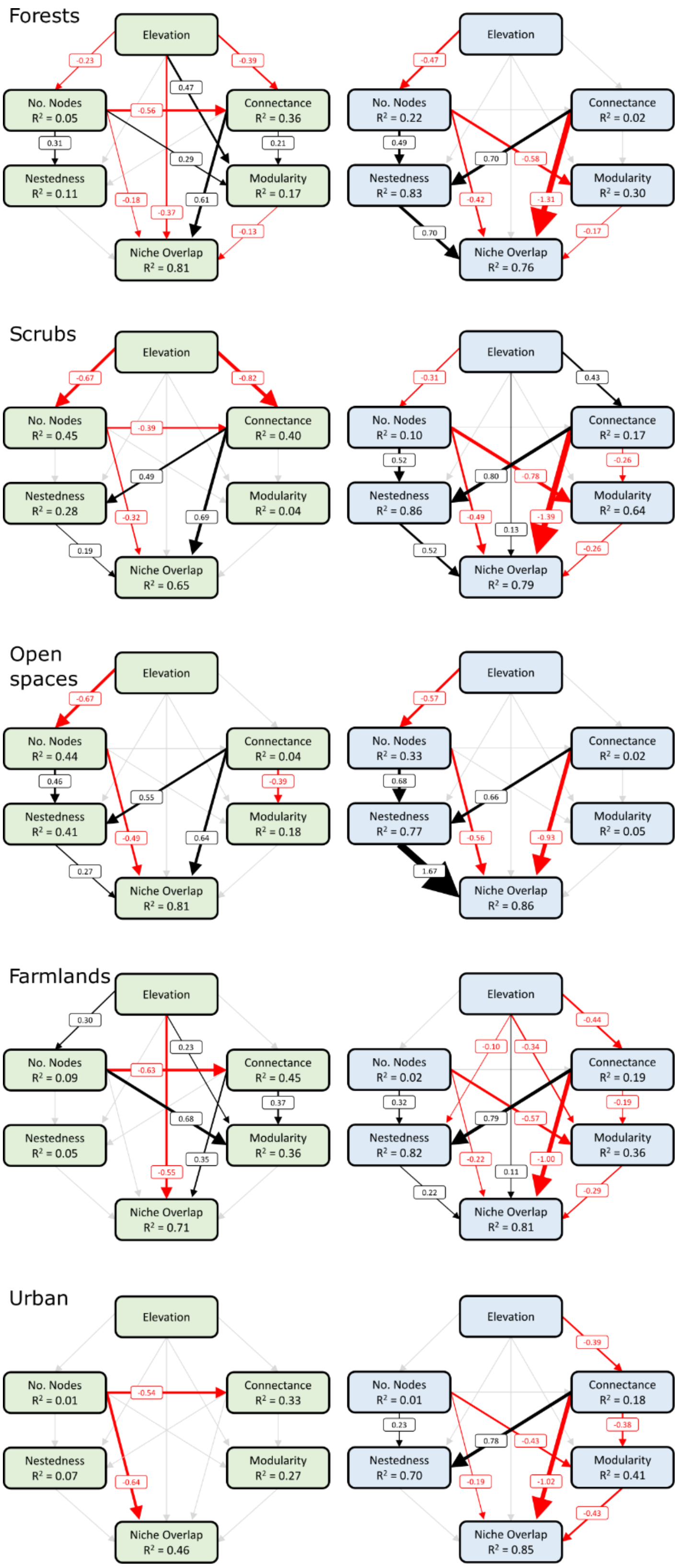


**Figure S1.** Piecewise structural equation modelling (SEM) analyses as of the main text, but with subsetted food webs of each dominant land-use type. We note that most of the detected causalities are subsets of the overall pattern (main text Figs. 3B–C), and farmlands exhibit significant yet opposite elevation to modularity and elevation to niche overlap influences in both green and blue food webs.

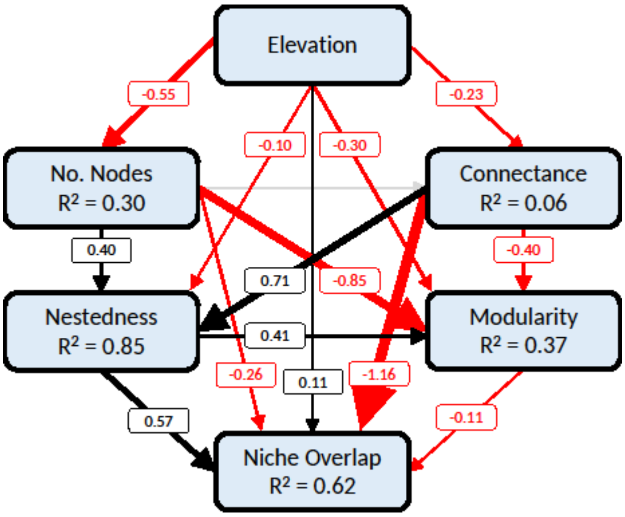


**Figure S2.** Piecewise structural equation modelling (SEM) analyses of the blue food webs as of the main text, but with a direct path from nestedness to modularity, which is unspecified in our model structure (main text Fig. 3B). The analysis suggests that adding such a path better explains the data. We note that with or without this unspecified path, all other detected causalities remain qualitatively and quantitatively consistent (main text Fig. 3C). Given such robustness, we present the results without this path in the main text, sticking with our literature-based model structure.

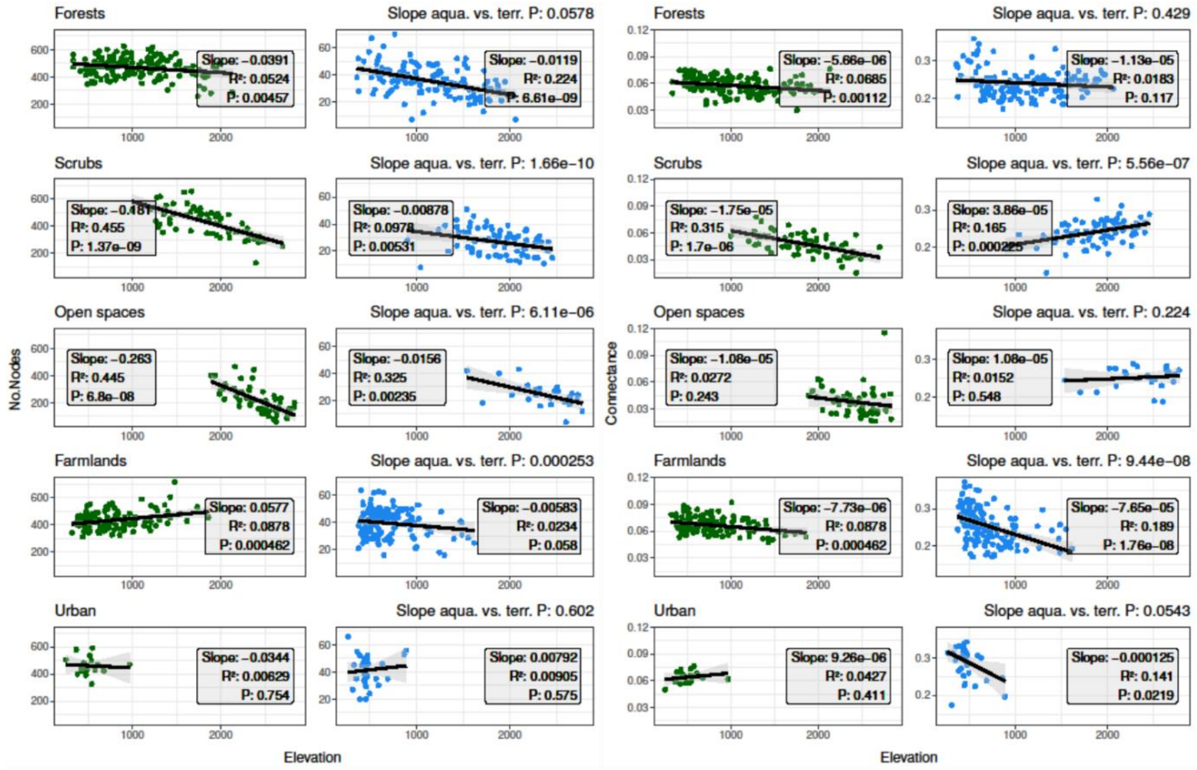


**Figure S3.** Detailed slope comparisons with linear models testing the effects of elevation on number of nodes (left panel) and connectance (right panel) with subsetted inferred food webs of each dominant land-use type. Overlaying theses land-type specific plots gives the scatterplots in main text Fig. 4, while the slope comparison stats here are summarised in the barplots there.

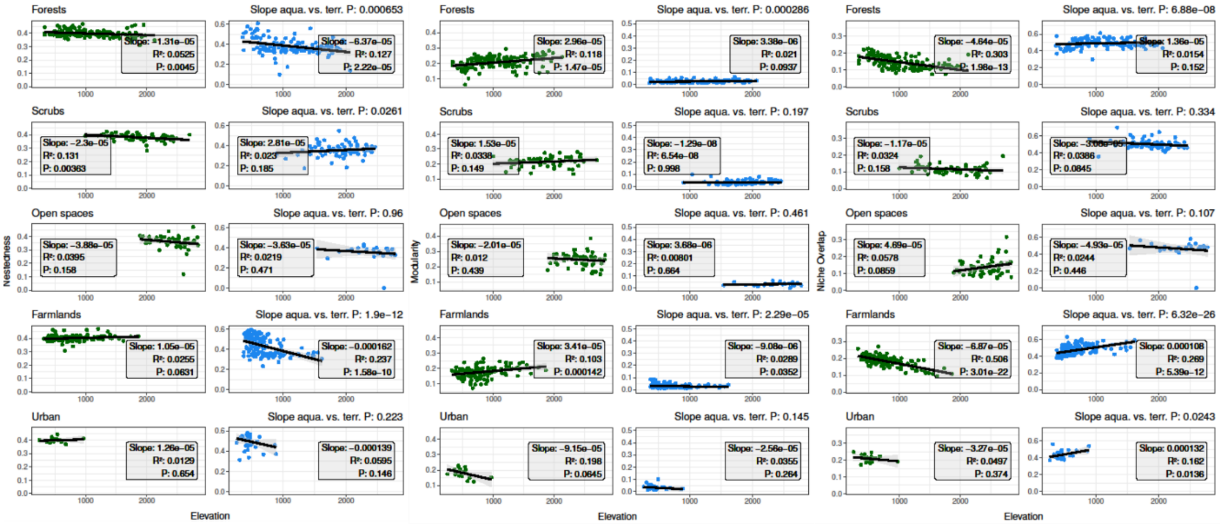


**Figure S4.** Detailed slope comparisons with linear models testing the effects of elevation on nestedness (left panel), modularity (middle panel), and consumers' niche overlap (right panel) with subsetted inferred food webs of each dominant land-use type. Overlaying theses land-type specific plots gives the scatterplots in main text Fig. 5, while the slope comparison stats here are summarised in the barplots there.

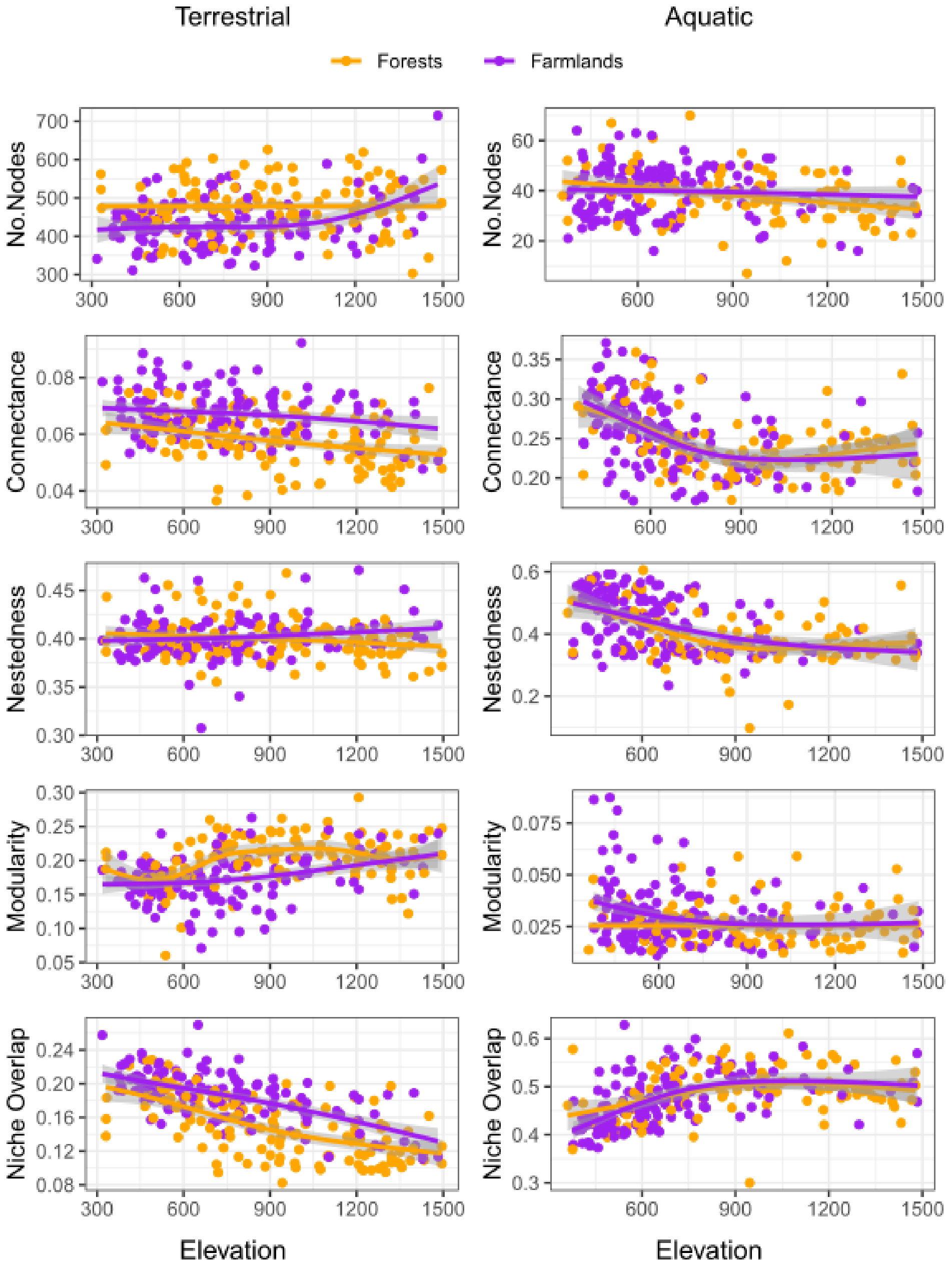


**Figure S5.** Comparisons with generalised additive models testing the effects of elevation on all food-web properties among subsetted food webs in forests versus farmlands, underneath 1500 m elevation. These two land-use types overlap their distributions in such an elevational segment and are thus comparable within. Solid lines and corresponding shades are the fitted regression and 95% CI, respectively.

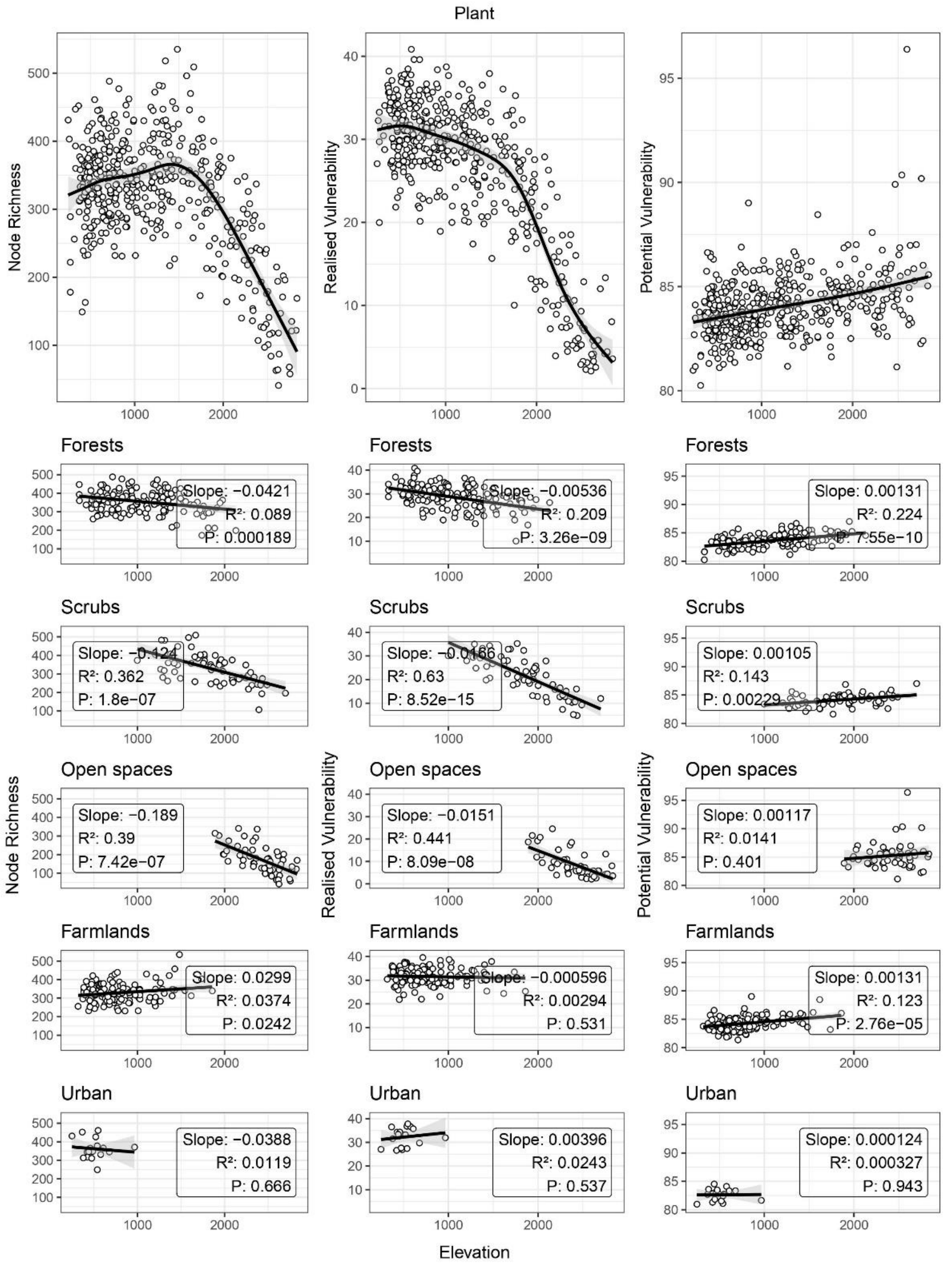


**Figure S6.** Node richness (left panel), realised vulnerability (middle panel), and potential vulnerability (right panel) of plants in assembled green food webs along elevation. The realised vulnerability is how many consumers feed on each plant in an inferred food web (based on consumers' occurrence at each site), whereas the potential one is the same measure in the metaweb (regional integration of trophic interactions). Each dot represents the mean value of an inferred food web. The top plot shows the overall pattern, whereas the below are patterns partitioned based on each dominant land-use type. The black lines are the fitted regression of generalised additive (overall) or linear (land-type specific) models with the corresponding shades the 95% CI.

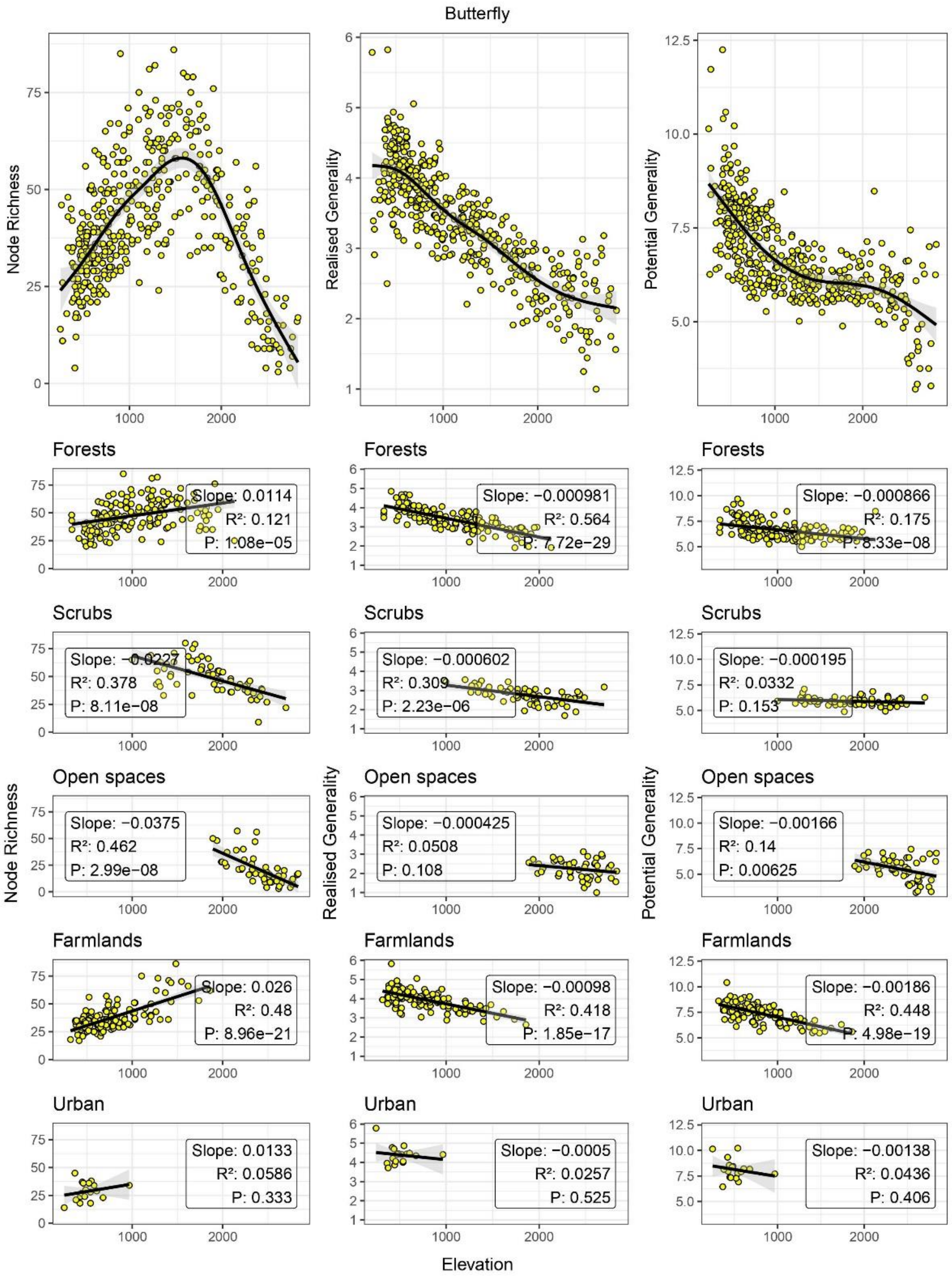


**Figure S7.** Node richness (left panel), realised generality (middle panel), and potential generality (right panel) of butterfly larva in assembled green food webs along elevation. The realised generality is the number of resources (host plants) that each focal butterfly feeds on in an inferred food web (based on resources' occurrence at each site), whereas the potential one is the same measure in the metaweb (regional integration of trophic interactions, i.e., more its biological diet breadth). Each dot represents the mean value of an inferred food web. The top plot shows the overall pattern, whereas the below are patterns partitioned based on each dominant land-use type. The black lines are the fitted regression of generalised additive (overall) or linear (land-type specific) models with the corresponding shades the 95% CI.

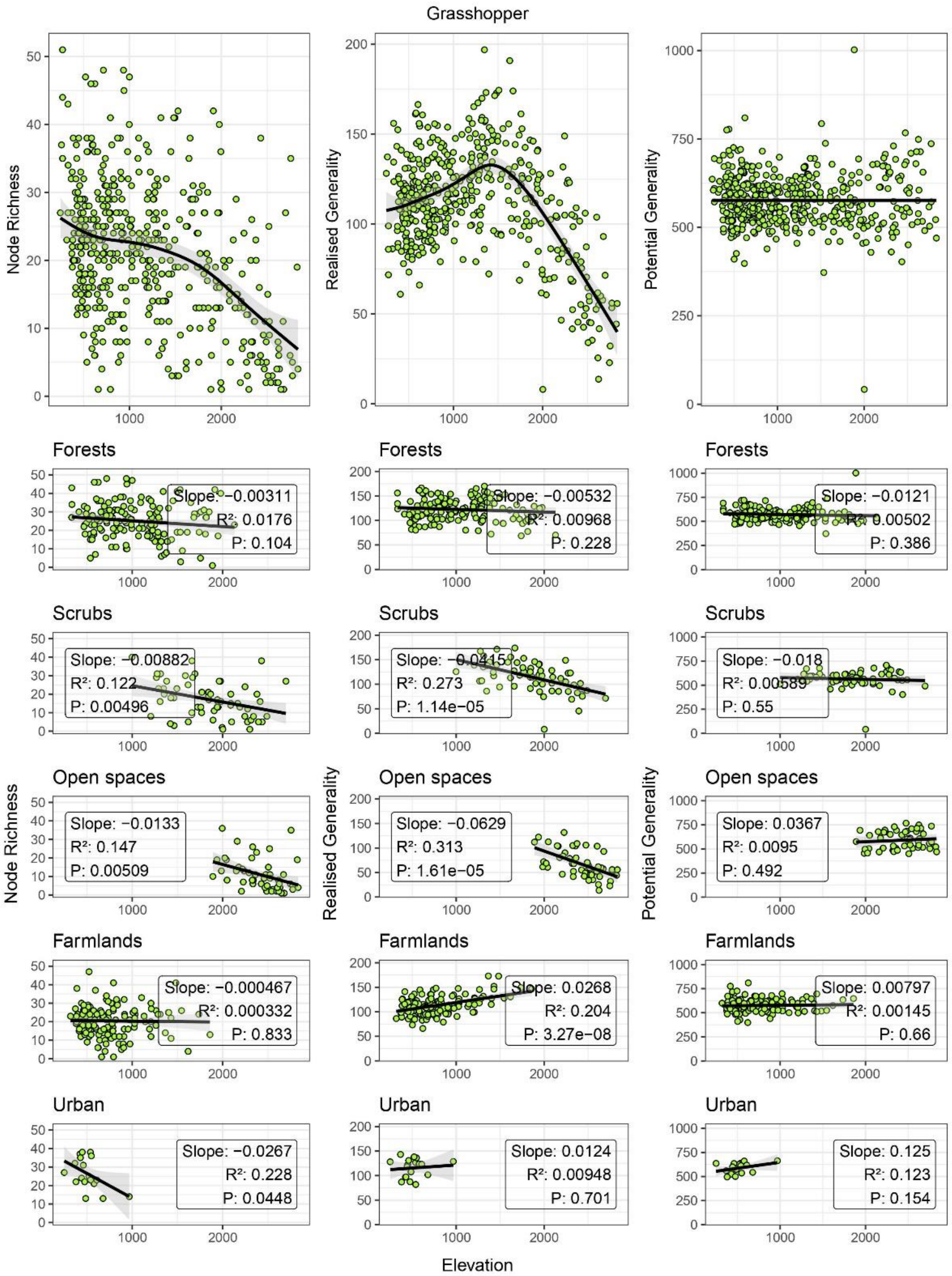


**Figure S8.** Node richness (left panel), realised generality (middle panel), and potential generality (right panel) of grasshoppers in assembled green food webs along elevation. The realised generality is the number of resources that each focal grasshopper feeds on in an inferred food web (based on resources' occurrence at each site), whereas the potential one is the same measure in the metaweb (regional integration of trophic interactions, i.e., more its biological diet breadth). Each dot represents the mean value of an inferred food web. The top plot shows the overall pattern, whereas the below are patterns partitioned based on each dominant land-use type. The black lines are the fitted regression of generalised additive (overall) or linear (land-type specific) models with the corresponding shades the 95% CI.

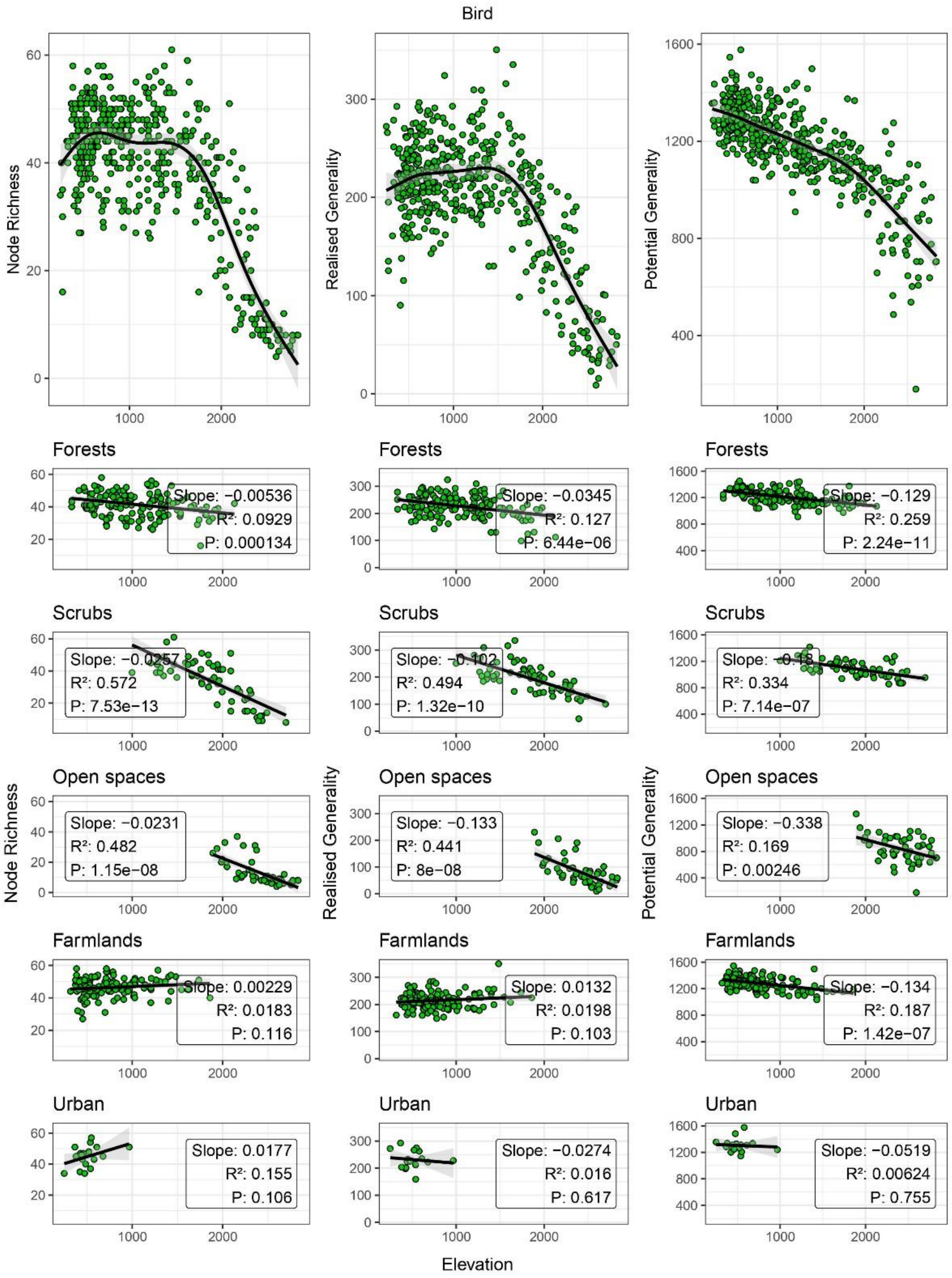


**Figure S9.** Node richness (left panel), realised generality (middle panel), and potential generality (right panel) of birds in assembled green food webs along elevation. The realised generality is the number of resources that each focal bird feeds on in an inferred food web (based on resources' occurrence at each site), whereas the potential one is the same measure in the metaweb (regional integration of trophic interactions, i.e., more its biological diet breadth). Each dot represents the mean value of an inferred food web. The top plot shows the overall pattern, whereas the below are patterns partitioned based on each dominant land-use type. The black lines are the fitted regression of generalised additive (overall) or linear (land-type specific) models with the corresponding shades the 95% CI.

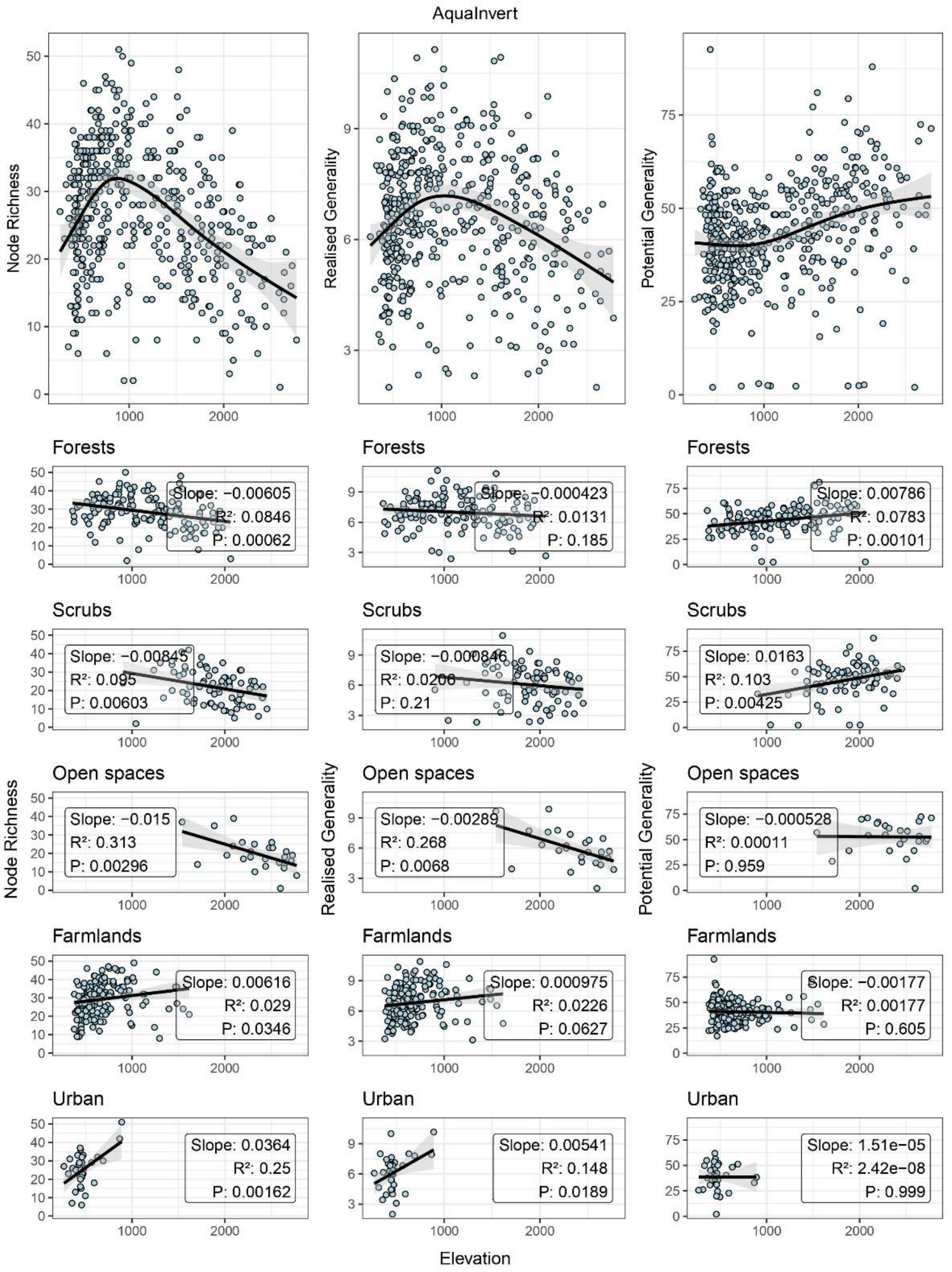


**Figure S10.** Node richness (left panel), realised generality (middle panel), and potential generality (right panel) of aquatic invertebrates in assembled blue food webs along elevation. The realised generality is the number of resources that each focal invertebrate feeds on in an inferred food web (based on resources' occurrence at each site), whereas the potential one is the same measure in the metaweb (regional integration of trophic interactions, i.e., more its biological diet breadth). Each dot represents the mean value of an inferred food web. The top plot shows the overall pattern, whereas the below are patterns partitioned based on each dominant land-use type. The black lines are the fitted regression of generalised additive (overall) or linear (land-type specific) models with the corresponding shades the 95% CI.

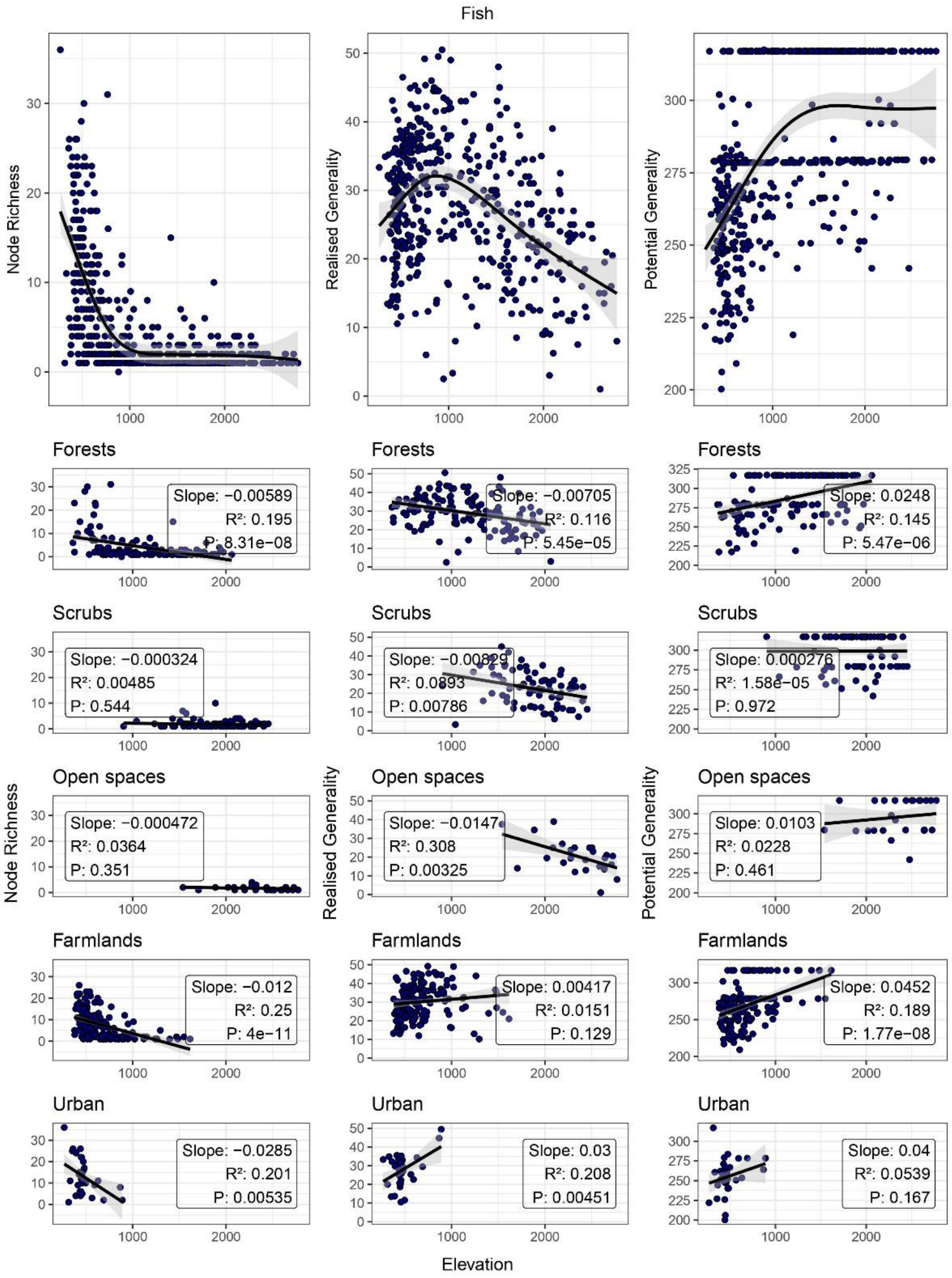


**Figure S11.** Node richness (left panel), realised generality (middle panel), and potential generality (right panel) of fishes in assembled blue food webs along elevation. The realised generality is the number of resources that each focal fish feeds on in an inferred food web (based on resources' occurrence at each site), whereas the potential one is the same measure in the metaweb (regional integration of trophic interactions, i.e., more its biological diet breadth). Each dot represents the mean value of an inferred food web. The top plot shows the overall pattern, whereas the below are patterns partitioned based on each dominant land-use type. The black lines are the fitted regression of generalised additive (overall) or linear (land-type specific) models with the corresponding shades the 95% CI. Note that our 5×5 $km^2$ fish occurrence resolution and 1×1 $km^2$ grid-averaged elevation may assign more fish species than actual occurring to some sites, particularly high-elevation ones.
